# Genomic surveillance enables suitability assessment of *Salmonella* gene targets used for culture-independent diagnostic testing

**DOI:** 10.1101/2020.01.07.898015

**Authors:** Rebecca J. Rockett, Alicia Arnott, Qinning Wang, Peter Howard, Vitali Sintchenko

## Abstract

*Salmonella* is a highly diverse genus consisting of over 2600 serovars responsible for high-burden food- and water-borne gastroenteritis worldwide. Sensitivity and specificity of PCR-based culture-independent diagnostic testing (CIDT) systems for *Salmonella*, which depend on a highly conserved gene target, can be affected by single nucleotide polymorphisms (SNPs), indels and genomic rearrangements within primer and probe sequences. This report demonstrates the value of prospectively collected genomic data for verifying CIDT targets.

We utilised the genomes of 3165 *Salmonella* isolates prospectively collected and sequenced in Australia. The sequence of *Salmonella* CIDT PCR gene targets (*ttrA, spaO* and *invA*) were systematically interrogated to measure nucleotide dissimilarity. Analysis of 52 different serovars and 79 MLST types demonstrated dissimilarity within and between PCR gene targets ranging between 0 – 81.3 SNP/Kbp (0 and 141 SNPs). Lowest average dissimilarity was observed in the *ttrA* target gene used by the Roche LightMix at 2.0 SNP/Kbp [range 0 – 46.7]), however entropy across the gene demonstrates it may not be the most stable CIDT target.

While debate continues over the benefits and pitfalls of replacing bacterial culture with molecular assays, the growing volumes of genomic surveillance data enable periodic regional reassessment and validation of CIDT targets against both prevalent and emerging serovars. If PCR systems are to become the primary screening and diagnostic tool for laboratory diagnosis of salmonellosis, ongoing monitoring of the genomic diversity in PCR target regions is warranted as is the potential inclusion of two *Salmonella* PCR targets into frontline diagnostic systems.

## Introduction

*Salmonella* is the leading cause of foodborne gastroenteritis worldwide, resulting in significant morbidity, mortality and economic cost.^1^ *Salmonella* is a highly diverse genus of zoonotic organisms, serologically classified into over 2600 serovars. The *Salmonella* genus is divided into two major species, *Salmonella enterica* and *Salmonella bongori. S. enterica* is considered the most pathogenic as the genome contains both *Salmonella* Pathogenicity Islands 1 and 2 (SPI1 and SPI2). *S. enterica* is further differentiated into six subspecies: *enterica* (Subspecies I), *salamae* (II), *arizonae* (IIIa), *diarizonae* (IIIb), *houtenae* (IV), and *indica* (VI). Non-typhoidal *Salmonella* (NTS) infections are primarily caused by serovars within Subspecies I, with *S. enterica* subspecies *enterica* serovar Typhimurium (STM) being a leading cause of foodborne outbreaks in many countries, including Australia. However, over 200 other serovars have been also reported as causal agents of NTS, including some that are unique to Australia.^2,3^ Prevalence of *Salmonella* serovars differs significantly between continents. For example, *S. enterica* subspecies *enterica* serovar Enteritidis is the most commonly detected serovar in Europe and North America, whereas STM predominates in Australia and New Zealand. ^4–7^

The public health, food safety and trade implications of foodborne *Salmonella* are such that a highly sensitive and specific surveillance system is required to ensure rapid detection and characterisation of foodborne outbreaks. The introduction and often complete replacement of traditional culture for *Salmonella* with culture-independent diagnostic testing (CIDT) as the front-end method of detecting *Salmonella* in stool samples has presented several challenges to public health laboratory surveillance.^5,8–10^ The reliance of all current CIDT platforms on a single gene target for the detection of a diverse pathogen such as *Salmonella* has raised important questions regarding ability of these CIDTs to detect uncommon serovars or emerging variants. As CIDTs become more widely used for *Salmonella* diagnosis, they become, in some cases, the only signal of a potential outbreak and the need for reflex culture. Therefore, any loss in CIDT sensitivity can have serious consequences for the diagnosis of individual cases as well as recognition and management of outbreaks.

Although PCR based techniques are considered extremely sensitive, rapid evolution within PCR targets has been documented for other pathogens. Mutations or indels can occur within CIDT primer and probe target regions and depending on the location of these mutations, CIDT sensitivity can be diminished and, in some cases, result in false negative results. For example, the sole reliance on CIDT to diagnose the highly conserved bacterium *Chlamydia trachomatis* is an example of the shortcomings of CIDT only testing algorithms.^11^ In 2006, it was reported that a *C. trachomatis* variant was circulating and this variant harboured a deletion in the PCR primer target region contained within the cryptic plasmid.^11^ This highly conserved multi-copy plasmid had been thought to be the ideal diagnostic target as it offered increased sensitivity for detecting *C. trachomatis.* However, emergence of a variant harbouring a deletion within the cryptic plasmid caused a complete loss of sensitivity in the majority of commercial CIDTs, resulting in the ongoing transmission of *C. trachomatis* and an increased number of severe sequalae after unrecognised and prolonged *C. trachomatis* infections.^12^ These false negative results for patients with variant *C. trachomatis* infections were only noted after a significant decrease in the incidence of *C. trachomatis* triggered an investigation by public health authorities. After the variant was identified it was found to be circulating for numerous years in sexual networks across the world. The response to this CIDT system failure was to design an additional *C. trachomatis* PCR target. The vulnerability of CIDT to generate false negative results due to nucleotide dissimilarity has led to the recommendation that a two-target system is needed for all frontline infectious disease CIDT assays.^13^ Despite this the current *Salmonella* CIDTs are often reliant on a single PCR target region.

Whole genome sequencing (WGS) provides the ultimate resolution to correctly identify outbreak clusters and detect food or environmental pathogen reservoirs.^14,15^ This ability triggered the rapid uptake of WGS as the preferred approach for public health surveillance of salmonellosis resulting in accumulation of genome sequence data. This genomic data provides an opportunity to examine variability of molecular targets employed by different CIDT platforms in the context of locally circulating *Salmonella* serovars. In this study we examined the variability in CIDT targets and the robustness of current CIDTs systems utilizing a comprehensive *Salmonella* genome collection spanning more than two summer seasons in New South Wales (NSW), Australia.

## Materials and Methods

### Isolate collections

*Salmonella* isolates included in this study represent all isolates referred to the NSW Enteric Reference Laboratory, ICPMR, NSW Health Pathology that underwent whole genome sequencing as part of routine public health outbreak investigations in NSW, Australia between October 2015 and December 2018 (n=3256). In addition, WGS was performed on 43 isolates collected from historical outbreaks of Salmonellosis where the causal *Salmonella* serovar had previously been described as native to Australia.^3^

### Nucleic acid extraction and library preparation

A single colony of overnight culture was used for DNA extraction and performed using the Geneaid Presto gDNA bacteria kit (Geneaid, Taiwan) as per the manufacturer’s instructions for Gram negative bacteria. DNA extracts were treated with 1 Unit of RNase. DNA libraries were prepared with the Nextera XT Library Preparation Kit, using 1ng of DNA in accordance with manufacturer’s instructions. Multiplexed libraries were sequenced using paired-end 150-bp chemistry on the NextSeq 500 (Illumina, Australia).

### Bioinformatic analysis of sequenced genomes

De-multiplexed sequencing reads with >1×10^7^ reads per isolate were trimmed^16^, based on a minimum quality read score of 20, then *de novo* assembled using SPAdes (version 3.13.0).^17^ The quality of *de novo* assemblies was assessed with Quast (5.0.2)^18^, only assemblies with <200 contigs and N50 > 50,000 bp were included in further analysis. MLST and serovar were inferred from final contigs using the *Salmonella enterica* pubMLST scheme (https://github.com/tseemann/mlst) and SISTR.^19^ The contigs were annotated with Prokka (version 1.13.3).^20^ Core genome analysis was conducted using Roary: the pan genome pipeline (version 3.12.0).^21^ Maximum-likelihood phylogeny of the pangenome was generated using the General Time Reversable (GTR+R4) model with IQ-TREE (version 1.6.3).^22^ Phylogeny and metadata were visualised with Microreact^23^ and Inkscape (https://inkscape.org/). CIDT gene targets *inv*A, *spa*O and *ttr*A were extracted from contigs using in-house Perl scripts and BLAST+.^24^ CIDT target genes from the closed reference genome *Salmonella enterica* subsp. *enterica* serovar Typhimurium str. LT2 (GeneBank Accession AE006468.2) were used as a reference. All CIDT nucleotide sequences extracted from isolate contigs were aligned to the reference using MAFFT.^25^ SNPs were called by their comparison to the reference genome using SNP-sites.^26^ SNP differences and the length of PCR target genes were used to calculate the dissimilarity index (SNP/Kbp) for each gene, however truncations were not included in SNP/Kbp calculations (gene lengths: *invA* 2065 bp, *spaO* 912 bp and *ttrA* 3063 bp). Entropy at each position within PCR target genes was measured from the PCR gene target alignments using the formula *H(l)* = Σ*f(b,l)*log_(base 2)_ *f(b,l)*. The core gene sequences including the gene targets *ttrA, spaO*, and *invA* for all 3165 isolates were deposited in the NCBI (BioProject PRJNA596817) (Supplemental Table 1).

### Primer and probe design and assessment

High homology regions of the PCR target genes (*ttrA, spaO*, and *invA)* were extracted from reference genomes of ten common causal serovars of Salmonellosis; Typhimurium, Enteritidis, Newport, Saintpaul, Virchow, Infantis, Hedelberg, Montevideo, Javiana, Muenchen and Braenderup. The reference *Salmonella enterica* genomes for these serovars were downloaded and the target gene sequences were extracted as outlined in the bioinformatic analyses described above (NCBI accession numbers for reference genomes: AE006468.2, MATG01.1, NC_011080.1, NC_011083.1, NC_-11294.1, NC_020307.1, NZ_CP007530.1, NZ_CP017727.1, NZ_CP022490.1, NZ_CP025094.1 and NZ_LN649235.1). Regions of homology between all sequences >300bp in length were employed to find the best primer and probe targets using the PrimerQuest Tool, Integrated DNA Technologies (https://sg.idtdna.com/Primerquest). The predicted oligonucleotides were then extracted from the genomes used in this study to investigate polymorphisms within the primer and probe sequences. The impact of any polymorphisms detected was based on both the number of SNPs in each oligonucleotide and the position of the SNPs within the oligonucleotide. A single SNP identified within the last 5 bp of the 3’end of the oligonucleotide were considered to have a moderate effect on PCR sensitivity, as were more than two SNPs within the same oligonucleotide. Significant sensitivity losses are predicted to occur when more than three SNPs are detected within the same oligonucleotide or more than one SNP within the last 5 bp at the 3’end of each oligonucleotide. Limited sensitivity losses were predicted for single SNPs outside the last 5 bp at the 3’end of the primer or probe.^27^

### Statistical Analysis

The diversity of NTS serovars in Australia between 2009 and 2017 was investigated using the National Notifiable Diseases Surveillance System Public *Salmonella* dataset (http://www9.health.gov.au/cda/source/). The total number of NTS serovars reported and case notifications of salmonellosis each year were used to calculate a Simpson’s index of diversity.

## Results

### Phylogenomic comparison of Salmonella genomes

A total of 3165 *Salmonella* isolates were included in this study. Isolate genomes that failed to meet sequence quality metrics or were sequenced multiple times were removed (n=134). The core genome shared by study isolates consisted of 3270 genes with a total length of 2,963,964 bp. Maximum likelihood core genome phylogeny was constructed using 98,254 polymorphic sites (median SNP distance 13,588 range 0 – 98,254) within core genes and demonstrated branching largely corresponding to serovar differentiation (Figure 1). The core genome included the PCR gene targets investigated in this study: *ttrA, spaO*, and *invA.* Assembly statistics for *Salmonella* genomes included in the study are outlined in Supplemental Table S1. The genomes investigated represented 52 different serovars and 79 MLST types, predominately Typhimurium (n=2174, 69%), including the monophasic Typhimurium variant I 4,[5],12:i:- (n=212, 7%), followed by Enteriditis (n=406, 13%) and Saintpaul (n=97, 3%). A detailed list of serovar and MLST types of all isolates is presented in Supplemental Table S2.

**Figure 1.**
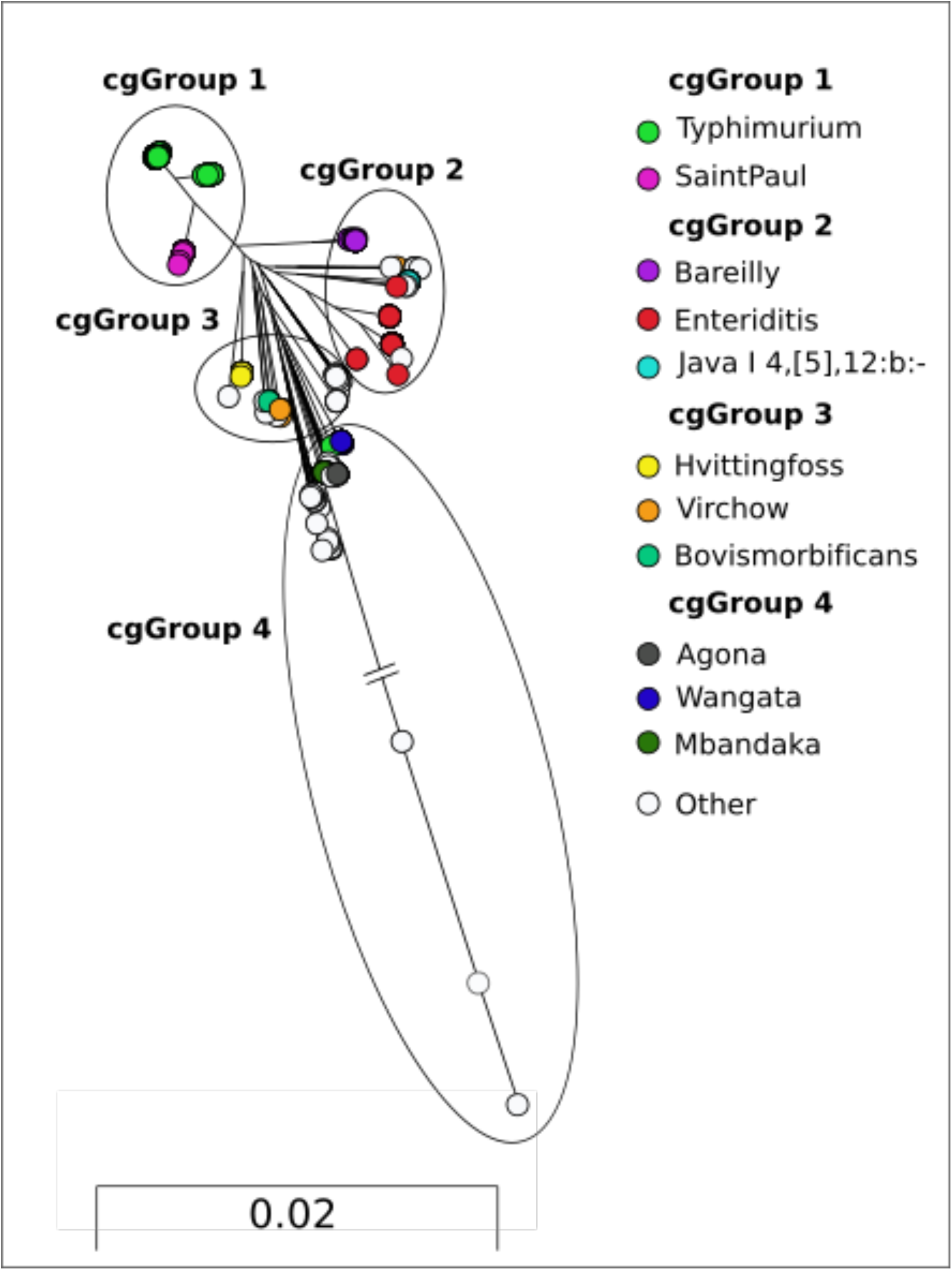
Maximum likelihood phylogeny constructed from 3270 core *Salmonella* genes. Coloured nodes represent serovars which contain more than 20 isolates, white nodes represent serovars with less than 20 isolates within the phylogeny. Long branches of cgGroup 4 have been truncated to aid visualisation.

To examine CIDT target nucleotide diversity, we divided the genomes into four groups (cgGroup 1-4) according to the phylogenetic analysis (Figure 1), with the aim of determining if CIDT target diversity correlated with core genome diversity. cgGroup 1 contained 2271 genomes and was dominated by the most common serovar in Australia, *Salmonella* Typhimurium (including monophasic S. Typhimurium; n=2174), in addition to serovar SaintPaul (n=97). cgGroup 2 consisted primarily of 515 genomes including serovars Enteritidis (n= 406), Bareilly (n=46) and Java (antigenic structure I 4, [5],12:b:-) (n=53). cgGroup 3 included 183 genomes and comprised primarily of serovars Hvittingfoss (n=106), Virchow (n=36) and Bovismorbificans (n=17). cgGroup 4 contained 196 genomes representing largely serovars Wangata (n=90), Mbandaka (n=40) and Agona (n=23). Long branches manifested in cgGroup 4 were attributed to the genomes of *S. enterica* subspecies *houtenae, diarizonae* and *salamae* (Figure 1). The branch lengths have been truncated to ease illustration of all core genome groups, the original tree can be found in Supplemental Figure 1. Notably, serovars that are unique to Australia, such as Adelaide, Chester, Orion and Tennessee, were also included in this cgGroup 4. cgGroups and their corresponding serovars and MLST profile is also presented in Supplemental Table S2.

### Nucleotide dissimilarity of PCR target genes

Significant nucleotide diversity was identified within all PCR target regions (Figure 2). Dissimilarity of CIDT targets within the same serovar was low. For example, the median dissimilarity indexes between only *S*. Typhimurium isolates within *invA, ttrA* and *spaO* nucleotide sequences in cgGroup 1 was 0.27 (range 0 – 1) for *invA*, 0 (range 0 – 10.8) for *ttrA* and 1.1 (range 0 – 1.1) for *spaO*, respectively. In contrast, significant dissimilarity was noted between other cgGroups containing a higher diversity of serovars. For *inv*A region, the median nucleotide dissimilarity indexes ranged from 6.8 (3.9 – 9.7), 3.9 (3.9 – 9.2) and 8.2 (0 - 51.3) for cgGroup 2, 3 and 4, respectively; for *ttr*A dissimilarity indexes were 5.2 (3.9 – 10.5), 5.2 (2 – 13.1), and 9.1 (6.5 – 46.1); and for *spa*O they ranged from 10.9 (1.1 – 12.1), 9.9 (1.1 – 13.2) and 8.6 (0 - 81.3), respectively. The lowest median dissimilarity between all *Salmonella* genomes was observed within the *ttrA* gene. Interestingly, the highest dissimilarity indexes were observed within genes from the *Salmonella* genomes in cgGroup 4 which included a representative from each of the *S. enterica* subspecies II to IV *salamae, diarizonae* and *houtenae*.

**Figure 2.**
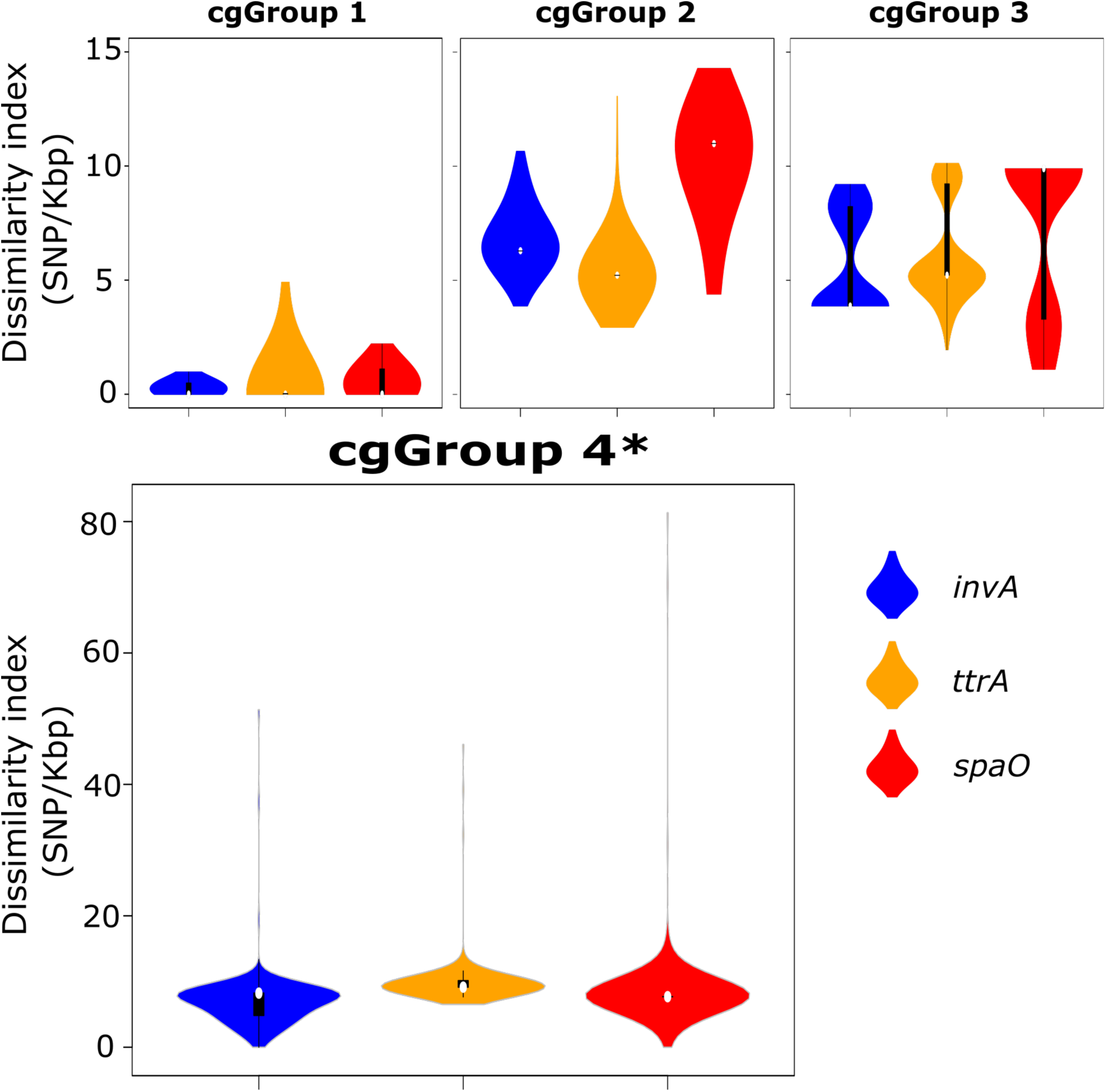
Nucleotide dissimilarity of CIDT PCR targets within each of the four phylogenetic groups. * Note increased dissimilarity scale for cgGroup4 genomes.

### Entropy of CIDT target genes

Entropy of CIDT target regions indicated that nucleotide dissimilarity was dispersed across the length of the sequence (Figure 3). Clear regions of conservation were seen in the 3’end of *invA* and 5’end of *spaO* genes sequences. However, polymorphisms were present throughout the *ttrA* nucleotide sequence. In addition, 16 of the genomes had truncations in 5’ and 3’ end of the *ttrA* nucleotide sequence with the length of truncation ranging from 60 to 3035 bp (median length 1337 bp). However, these same genomes had lower quality assembly metrics compared to the genomes from the entire collection. The medium number of contigs in the 16 assemblies that contained a truncated *ttrA* gene sequence was 99.5 (34 – 135) as compared to that of the entire genomes analysed at 78 (24-169). The median N50 value for genomes containing a truncated *ttrA* nucleotide sequence was slightly lower at 139,974 bp (66,538–261,187 bp) as compared to all genomes analysed at 172,206 bp (52,145 –518,076) (Supplemental Table S3). Therefore, these truncations may have resulted from *de novo* assembly errors.

**Figure 3.**
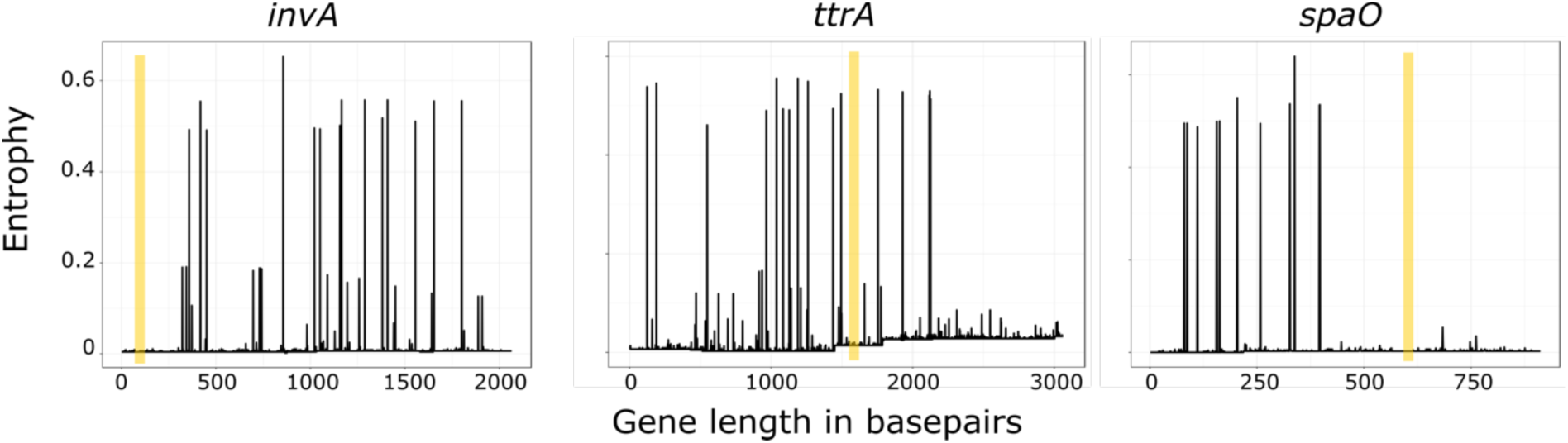
The entropy across the nucleotide sequence of the three CIDT targets, *invA, ttrA* and *spaO* genes. Yellow boxes highlight the optimal primer and probe target regions within each gene.

### Nucleotide diversity in primer and probe targets

*In silico* prediction of primer and probe targets for each gene was performed using regions of PCR target genes homologous amongst ten common *Salmonella enterica* serovars. Predicted gene sequences and oligonucleotide parameters are specified in Table 1. Entropy across the RT-PCR amplicon for each gene target is depicted in Figure 3. When comparing the entropy of the CIDT target across all genomes in this study, the RT-PCR target regions remain in areas of low nucleotide variability. The specific oligonucleotide sequences were extracted from each genome, with only 2% (60/3165) genomes contained SNPs within proposed oligonucleotide sequences. Four, 12 and 44 genomes contained oligonucleotide mutations in the *invA, ttrA* and *spaO* RT-PCR oligonucleotides, respectively. Most single SNPs in either the forward or reverse primers and probe oligonucleotide were expected to have a small impact on assay sensitivity (48/60). However, the remaining 12 genomes had multiple SNPs within a single oligonucleotide or contained a single mutation in multiple oligonucleotides. Multiple mutations in single oligonucleotides or mutations across all oligonucleotides of an RT-PCR are likely to lead to significant sensitivity losses and in some cases false negative results. The genomes harbouring multiple mutations (12/60) were all contained within cgGroup 4 which contained the most divergent core genomes (Table 2).

**Table 1.**
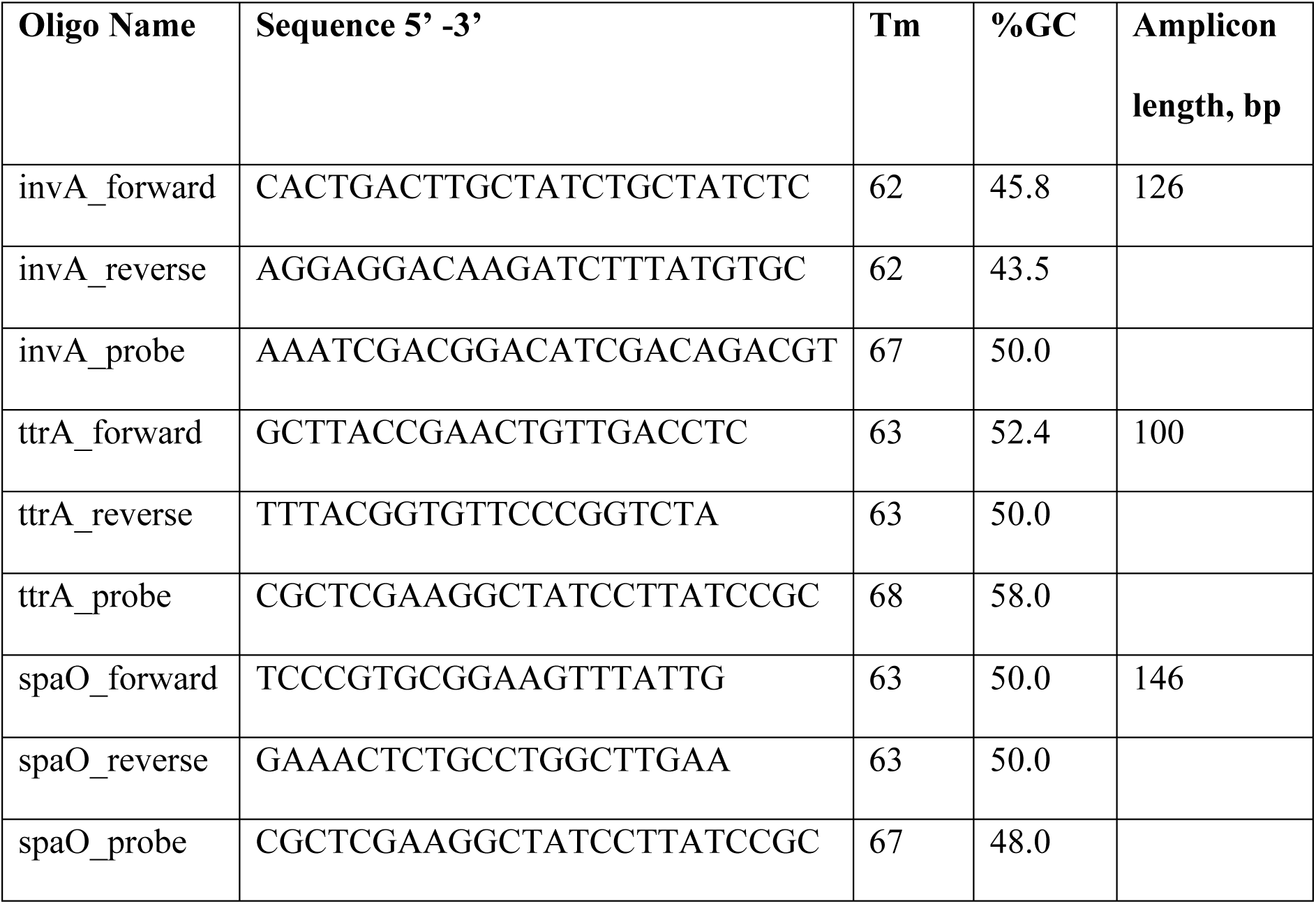
Oligonucleotide sequences and PCR parameters designed for this study.

**Table 2.** Polymorphisms in RT-PCR assay oligonucleotide sequences, with predicted loss in assay sensitivity.

| Gene | Oligo effected | Position in oligo from 5' end | Predicted loss in sensitivity | Serovar (total number of genomes affected encoding that serovar) | Total No. of genomes | Core genome Group |
| --- | --- | --- | --- | --- | --- | --- |
| ***invA*** | Forward primer | 16<sup>th</sup> base | moderate | IV 43:z4,z23:- (1), ST237 | 1 | 4 |
|  | Forward primer | 1<sup>st</sup> base | limited | II1,4,[5],12,[27] (1)* | 1 | 4 |
|  | Probe | 1<sup>st</sup> base | limited | Clanvillian (1)* | 1 | 4 |
|  | Reverse primer | 12<sup>th</sup> base | limited | IIIb,50:k:z35 (1)*<br>II1,4,[5],12,[27] (1)* | 2 | 4 |
| ***ttrA*** | Forward Primer | 5<sup>th</sup> base | limited | Enteriditis (469) | 1 | 2 |
|  | Forward Primer | 4<sup>th</sup> base | limited | IV 43:z4,z23:- (1), ST237 | 1 | 4 |
|  | Forward Primer | 6<sup>th</sup> base | limited | Senftenberg (2),<br>ST210 | 1 | 4 |
|  | Probe | 2 SNPs,<br>2<sup>nd</sup> and<br>11<sup>th</sup> base | moderate | Tennessee (3), ST319 | 3 | 4 |
|  | Probe | 11<sup>th</sup> base | limited | Havana (4), ST595,<br>ST578 | 3 | 4 |
|  | Probe | 20<sup>th</sup> base | moderate | Typhimurium (2175),<br>ST19 | 1 | 1 |
|  | Probe | 1<sup>st</sup> base | limited | IV 43:z4,z23:- (1), ST237 | 1 | 4 |
|  | Reverse primer | 14<sup>th</sup> base | limited | IV 43:z4,z23:- (1), ST237 | 1 | 4 |
| *spaO* | Forward primer | 4 and 12<sup>th</sup> base | moderate | IIIb,50:k:z35 (1)* | 1 | 4 |
|  | Reverse primer | 12<sup>th</sup> base | limited | Mbandaka (40), ST413 | 40 | 4 |
|  | Reverse primer | 4,12 and 15<sup>th</sup> base | significant | II1,4,[5],12,[27] (1)* | 1 | 4 |
|  | Reverse primer | 3, 5 and 15<sup>th</sup> base | significant | IV 43:z4,z23:- (1), ST237 | 1 | 4 |
|  | Reverse primer | 5 and 15<sup>th</sup> base | moderate | IIIb,50:k:z35 (1)* | 1 | 4 |

\* Novel allele combination

### Temporal trends in Australian non-typhoidal Salmonella serovar diversity

Based on the Australian National Notifiable Diseases Surveillance System Public *Salmonella* dataset, there were more than 200 serovars identified in association with more than 14,000 salmonellosis cases each year during the period of 2009 and 2017 in Australia. A gradual increase in serotype diversity and reported incidence of non-typhoidal salmonellosis was detected between 2009 and 2017 (Figure 4). CIDT for *Salmonella* diagnosis were introduced widely across Australia in late 2013, followed by notable increases in the Simpson’s diversity index after 2015. Subspecies II-IV account for around 1% of Salmonellosis cases reported each year in Australia.

**Figure 4.**
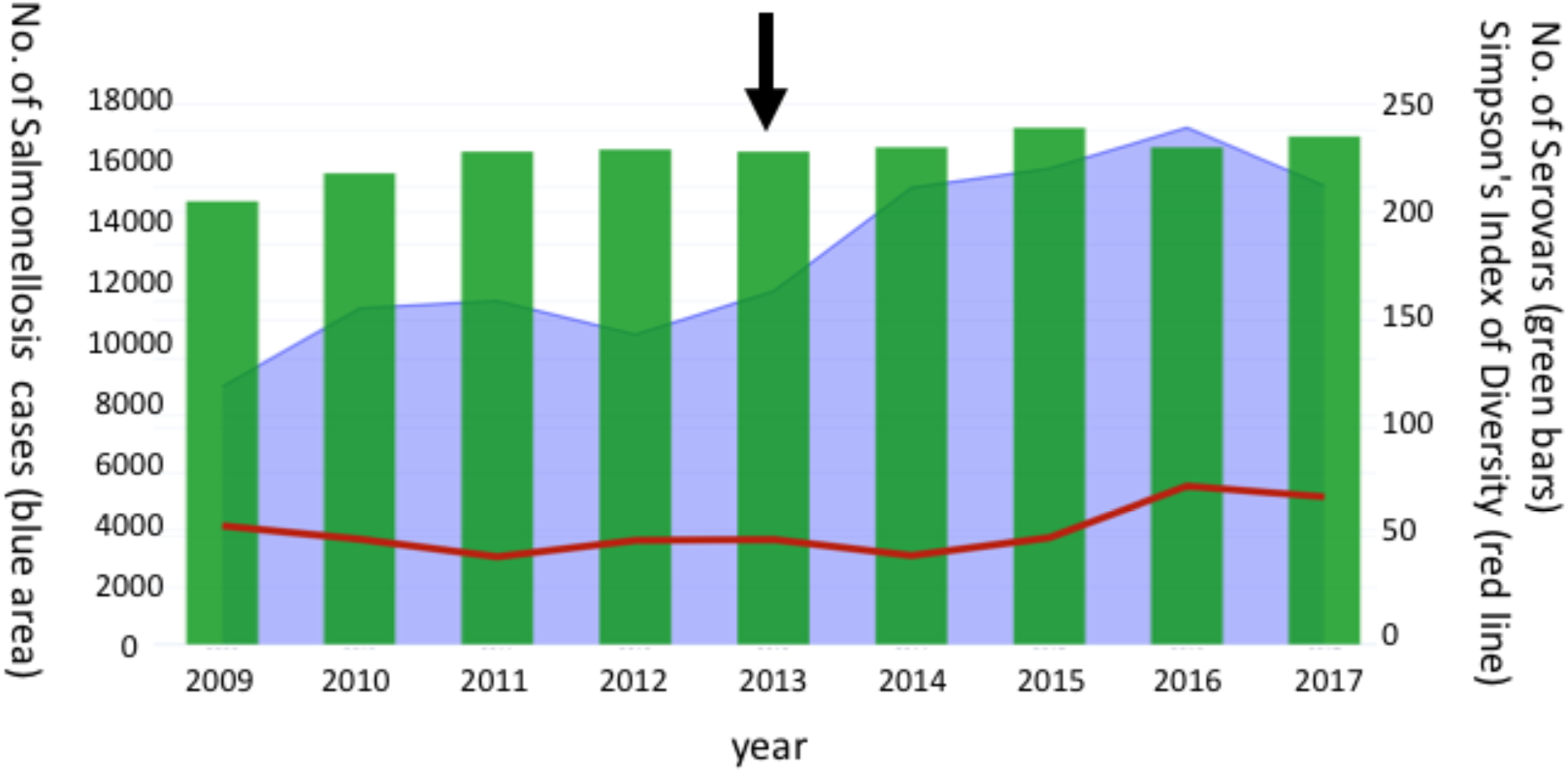
Serotype diversity of notified cases of Salmonellosis in Australia between 2009 and 2017. *CIDTs platforms became available in 2013 in Australia, indicated by the black arrow. Simpson’s index of Diversity was multiplied by a factor of 50 to aid visual representation on the graph above.

## Discussion

WGS is being increasingly utilized to investigate outbreaks of *Salmonella* and to accurately identify food and environmental sources of infection. This high-resolution approach has greatly enhanced the speed and accuracy of public health interventions ensuring food safety. Genome sequencing data collected as part of prospective public health surveillance has also enabled the ongoing sensitivity of foodborne outbreak surveillance to be determined and monitored, as described in the present study. Public health authorities have raised concerns that the increasing reliance on CIDTs as the only tool to detect pathogens responsible for foodborne disease will significantly reduce the sensitivity of public health surveillance systems as serotyping and WGS require salmonella to be isolated by solid medium culture.^8,9,28^ The replacement of conventional culture with CIDT reduces the number of isolates available for WGS and therefore the ability of public health laboratories to identify and control outbreaks of salmonellosis. In Australia, it has been reported that 12-18% of Salmonella positive stool samples are identified based solely on CIDT testing.^8,29^ In North America and Canada, Salmonella positive stool samples are reflectively cultured after a positive *Salmonella* CIDT at public health laboratories in an attempt to boost the number of Salmonella isolates available for WGS. Australia has not yet widely adopted reflex culture after a positive Salmonella CIDT however it may be required to maintain the current WGS based surveillance systems.^5,10^

The CIDT platforms offer highly automated ‘swab to result’ systems with high throughput, rapid turnaround time and reduced testing costs. However, most commercial assays offer a single PCR target for *Salmonella* detection, with the specific primer and probe target sequences difficult to uncover as they are commonly proprietary. In this study we systematically interrogated three genes used as PCR targets by different CIDT platforms including *ttrA* (LightMix Modular *Salmonella* CE-IVD, Roche/TIB MOLBIOL)^30^, *spaO* (BD Max)^31^ and *invA*^32^ (a common in-house *Salmonella* PCR target). While our genomic analysis based on WGS data allowed the assessment of dissimilarity of CIDT targets, the proprietary nature of PCR primers and probes made it difficult to accurately estimate the extent of the potential sensitivity losses. We were, however, reassured that significant changes in the spectrum and diversity of Salmonella serovars do not appear to have occurred despite the recent introduction of CIDT systems for detection of bacterial enteropathogens in stool samples from patients with gastroenteritis in Australia. The Australian CIDT market has been dominated by commercial systems, namely the BD Max Enteric Bacterial Panel PCR Assay and the Roche/TIB MOLBIOL LightMix Modular Gastro Bacteria Multiplex testing. The recent implementation of other panels, including the BioFire FilmArray GI panels for which the gene targets are not disclosed as well as non-commercial PCR CIDT, has only recently occurred within the Australian diagnostic testing landscape and hence their impact on *Salmonella* surveillance is yet to be determined.

Dissimilarity of PCR targets occurred within each of the 52 Salmonella serovars investigated; the highest levels of dissimilarity were noted amongst the three genomes falling outside of *S. enterica* subspecies enterica Subspecies I. Although *Salmonella* Subspecies I isolates predominate as the causal agent of foodborne outbreaks in Australia and elsewhere, it should be noted that current CIDT may have diminished sensitivity to detect the less common *S. enterica* subspecies with potential for emergence as human pathogens, especially isolates from predominantly zoonotic Subspecies IV and VI.^24^ Amongst the individual genes, entropy across PCR targets suggests that the *spaO* gene may contain a more conserved PCR target region, while other targets may be affected by dissimilarity across the entire gene. In particular, truncations of *ttrA* gene region were present in a small number of genomes. The *ttrA* gene encodes tetrathionate reductase Subunit A and is part of the ttrRSBCA operon required for tetrathionate respiration and located in close proximity to Salmonella Pathogenicity Island 2. The integration of virulence factors or other evolutionary pressures may increase the variability in this particular CIDT target, but these truncations will need to be validated in other datasets to exclude bioinformatic assembly errors.

Truncations aside, the effect of polymorphisms in oligonucleotides on assay sensitivity is difficult to assess and can be greatly affected by a number of parameters. Generally, the largest sensitivity losses are seen with mutations at the 3’end of primer sequences.^33^ However, the composition of the mismatched base also plays a role as purine-pyrimidine mismatches are generally less detrimental than purine-purine/pyrimidine-pyrimidine polymorphisms.^27,34,35^ More general considerations are also important, including the master mix composition and the number of multiplexed primer and probe combinations. Generally, the more primer pairs included in a multiplex reaction the greater the loss of sensitivity for the mismatched primer set, particularly in poly-microbial intestinal infections. These observations are especially relevant and important as prediction models indicate that even a small loss of PCR sensitivity could be detrimental to outbreak detection.

In conclusion, the growing volumes of genomic surveillance data allow ongoing local reassessment and validation of CIDT targets representing prevalent and emerging serotypes of public health significance. Nucleotide dissimilarity of CIDT targets in different serovars of *Salmonella* Subspecies IV and VI may affect the public health surveillance of non-typhoidal salmonellosis in endemic areas. If CIDT systems are to become the primary screening and diagnostic tool for laboratory diagnosis of salmonellosis, ongoing monitoring of the genomic diversity in PCR target regions is warranted. A two gene target detection system is also recommended and would limit the potential of false negative CIDT results for *Salmonella*.

## Acknowledgments

This work was supported by the Prevention Research Support Program, funded by the New South Wales Ministry of Health.

The authors would like to acknowledge Nathan Bachmann and Karen-Ann Gray for their assistance with perl scripting and manuscript editing. In addition, Carl Banting, Chayanika Biswas, Rajat Dhakal, Jenny Draper, Mailie Gall, Andrew Ginn, Aneliese Goodwin, Karen-Ann Gray, Connie Lam, Dean Jalocon, Elena Martinez, Ranjeeta Menon, Eby Sim, and Verlaine Timms at The Microbial Genomics Reference Laboratory, NSW Pathology for their assistance with genome sequencing and bioinformatic analysis.

The NSW Enteric Reference Laboratory, NSW Pathology for assistance with serotyping, identification and access to historical *Salmonella* collections. Rosemarie Sadsad and The Sydney Informatics Hub along with The University of Sydney High performance computing cluster for bioinformatic assistance and computing capacity.

## Supplementary Material

**Supplemental Table S1.** *de novo* assembly metrics of *Salmonella* genomes.

| *de novo* assembly metrics | Median (range) |
| --- | --- |
| Number of contigs (>1000bp) | 78 (24-169) |
| Genome size | 4,879,273 (4,185,374 - 5,283,669) |
| GC content (%) | 52.2 (51.5 - 54.5) |
| N50 | 172,206 (52,145 - 518,076) |
| N75 | 75607 (27,194 - 253,326) |

**Supplemental Table S2.** Summary of serovar, sequence type and cgGroup of all isolates included in the study.

| Serovar | Sequence Type (ST) | Total No. of isolates per serovar | Total No. of isolates per ST | cgGroup per ST |
| --- | --- | --- | --- | --- |
| **Aberdeen** |  | 1 |  |  |
|  | 426 |  | 1 | 3 |
| **Abortusovis** |  | 4 |  |  |
|  | 768 |  | 4 | 2 |
| **Adelaide** |  | 4 |  |  |
|  | 440 |  | 4 | 4 |
| **Agona** |  | 23 |  |  |
|  | 13 |  | 23 | 4 |
| **Agoueve** |  | 1 |  |  |
|  | 286 |  | 1 | 4 |
| **Apeyeme** |  | 1 |  |  |
|  | 1546 |  | 1 | 4 |
| **Bahrenfeld** |  | 2 |  |  |
|  | 859 |  | 1 | 4 |
|  | - |  | 1 | 4 |
| **Bareilly** |  | 46 |  |  |
|  | 203 |  | 41 | 2 |
|  | 909 |  | 5 | 2 |
| **Birkenhead** |  | 1 |  |  |
|  | 424 |  | 1 | 3 |
| **Bovismorbificans** |  | 17 |  |  |
|  | 377 |  | 16 | 3 |
|  | 1499 |  | 1 | 3 |
| **Broughton** |  | 1 |  |  |
|  | - |  | 1 | 4 |
| **Cerro** |  | 1 |  |  |
|  | 3548 |  | 1 | 4 |
| **Chester** |  | 6 |  |  |
|  | 343 |  | 5 | 4 |
|  | - |  | 1 | 4 |
| **Clanvillian** |  | 1 |  |  |
|  | - |  | 1 | 2 |
| **Enteritidis** |  | 406 |  |  |
|  | 10 |  | 1 | 2 |
|  | 11 |  | 301 | 2 |
|  | 74 |  | 1 | 2 |
|  | 180 |  | 33 | 2 |
|  | 1925 |  | 58 | 2 |
|  | 1972 |  | 2 | 2 |
|  | 3233 |  | 4 | 2 |
|  | 3304 |  | 5 | 2 |
|  | - |  | 1 | 2 |
| **Galiema** |  | 1 |  |  |
|  | 601 |  | 1 | 3 |
| **Gallinarum** |  | 1 |  |  |
|  | 92 |  | 1 | 2 |
| **Gatineau \| Orion** |  | 5 |  |  |
|  | 580 |  | 3 | 4 |
|  | 639 |  | 2 | 4 |
| **Give** |  | 1 |  |  |
|  | 516 |  | 1 | 4 |
| **Havana** |  | 4 |  |  |
|  | 578 |  | 3 | 4 |
|  | 595 |  | 1 | 4 |
| **Hvittingfoss** |  | 107 |  |  |
|  | 434 |  | 94 | 3 |
|  | 446 |  | 3 | 3 |
|  | 2062 |  | 10 | 3 |
| **I 4,[5],12:b:-** |  | 53 |  |  |
|  | 2358 |  | 1 | 2 |
|  | 4959 |  | 52 | 2 |
| **II 1,4,[5],12,[27]:b:[e,n,x]** |  | 1 |  |  |
|  | 2116 |  | 1 | 4 |
| **IIIb 50:k:z35** |  | 1 |  |  |
|  | - |  | 1 | 4 |
| **Infantis** |  | 7 |  |  |
|  | 32 |  | 7 | 3 |
| **IV 43:z4,z23:-** |  | 1 |  |  |
|  | 237 |  | 1 | 4 |
| **Litchfield** |  | 1 |  |  |
|  | 4491 |  | 1 | 3 |
| **Manchester** |  | 1 |  |  |
|  | 491 |  | 1 | 3 |
| **Mbandaka** |  | 40 |  |  |
|  | 413 |  | 40 | 4 |
| **Mgulani** |  | 1 |  |  |
|  | 2033 |  | 1 | 3 |
| **Minnesota** |  | 1 |  |  |
|  | 548 |  | 1 | 4 |
| **Mishmarhaemek** |  | 1 |  |  |
|  | 2666 |  | 1 | 3 |
| **Montevideo** |  | 1 |  |  |
|  | - |  | 1 | 3 |
| **Muenchen** |  | 5 |  |  |
|  | 82 |  | 3 | 3/4 |
|  | 112 |  | 1 | 3 |
|  | 3211 |  | 1 | 3 |
| **Newport** |  | 1 |  |  |
|  | 31 |  | 1 | 3 |
| **Oesterbro** |  | 1 |  |  |
|  | 466 |  | 1 | 3 |
| **Oranienburg** |  | 1 |  |  |
|  | 23 |  | 1 | 4 |
| **Orientalis** |  | 1 |  |  |
|  | 558 |  | 1 | 4 |
| **Orion** |  | 1 |  |  |
|  | 639 |  | 1 | 4 |
| **Paratyphi A** |  | 1 |  |  |
|  | 129 |  | 1 | 4 |
| **Poona** |  | 1 |  |  |
|  | 1069 |  | 1 | 3 |
| **Potsdam** |  | 1 |  |  |
|  | 2462 |  | 1 | 2 |
| **Saintpaul** |  | 97 |  |  |
|  | 27 |  | 2 | 1 |
|  | 50 |  | 94 | 1 |
|  | 3528 |  | 1 | 1 |
| **Senftenberg** |  | 2 |  |  |
|  | 185 |  | 1 | 4 |
|  | 210 |  | 1 | 4 |
| **Singapore** |  | 2 |  |  |
|  | 462 |  | 1 | 3 |
|  | 501 |  | 1 | 4 |
| **Stanley** |  | 2 |  |  |
|  | 29 |  | 1 | 2 |
|  | 2299 |  | 1 | 2 |
| **Tennessee** |  | 3 |  |  |
|  | 319 |  | 3 | 4 |
| **Typhi** |  | 1 |  |  |
|  | 2 |  | 1 | 4 |
| **Typhimurium** |  | 1959 |  |  |
|  | 19 |  | 1881 | 1 |
|  | 34 |  | 6 | 1 |
|  | 36 |  | 31 | 1 |
|  | 313 |  | 1 | 1 |
|  | 513 |  | 1 | 4 |
|  | 2066 |  | 3 | 1 |
|  | 2089 |  | 5 | 1 |
|  | 2297 |  | 1 | 1 |
|  | - |  | 34 | 1 |
| **TyphimuriumI 4,[5],12:i:-** |  | 212 |  |  |
|  | 19 |  | 1 | 1 |
|  | 34 |  | 208 | 1 |
|  | - |  | 3 | 1 |
| **Virchow** |  | 37 |  |  |
|  | 16 |  | 35 | 3 |
|  | 197 |  | 1 | 2 |
|  | 359 |  | 1 | 3 |
| **Wangata** |  | 90 |  |  |
|  | 523 |  | 90 | 4 |
| **Grand Total** |  | **3165** | **3165** |  |

**Supplementary Figure 1.**
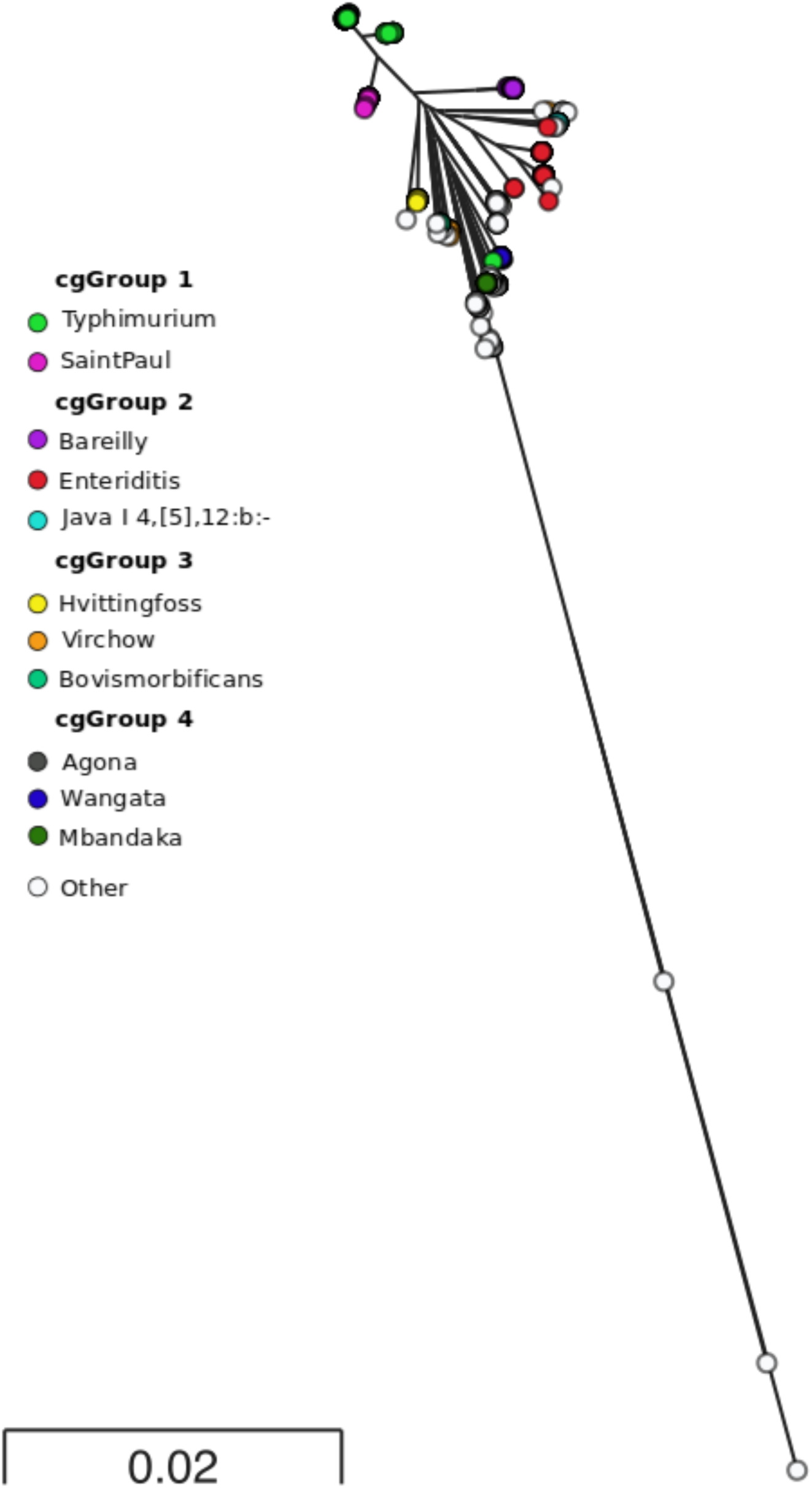
Maximum likelihood phylogeny constructed from 3270 core *Salmonella* genes. Coloured nodes represent serovars which contain more than 20 isolates, white nodes represent serovars with less than 20 isolates within the phylogeny.

